# Revealing compartmentalised membrane diffusion in living cells with interferometric scattering microscopy

**DOI:** 10.1101/091736

**Authors:** G. de Wit, D. Albrecht, H. Ewers, P. Kukura

## Abstract

Single-particle tracking is a powerful tool for studying single molecule behaviour involving plasma membrane-associated events in cells. Here, we show that interferometric scattering microscopy (iSCAT) combined with gold nanoparticle labeling can be used to follow the motion of membrane proteins in the plasma membrane of live cultured mammalian cell lines and hippocampal neurons. The unique combination of microsecond temporal resolution and nanometer spatial precision reveals signatures of a compartmentalised plasma membrane in neurons.

---

Single-particle tracking (SPT) generally involves tagging a single molecule with a probe, and detecting its position in a series of images, using computational image processing to achieve sub-pixel localisation precision.^1^ These trajectories contain information on the diffusion coefficient of the molecule, its momentum and possible underlying forces controlling its motion.^2-5^ Furthermore, the trajectory can be correlated to features of the cell or the motion of interacting partners, providing a means to analyse dynamic changes in the behaviour of single molecules. While single particle tracking was first established using gold beads^6^, current implementations also make use of quantum dots (QDs) or single fluorescent dyes as probes, which have lead to the development of high-density methods.^7-9^

The enormous advantage of fluorescence emission as a contrast mechanism in terms of maximal background suppression is somewhat offset by restrictions in terms of the achievable simultaneous imaging speed and precision. Scattering labels, such as gold nanoparticles, are not subject to the photophysical and photochemical limitations of fluorescent dyes and can thus in principle achieve much higher spatiotemporal precision. Gold particles have been used extensively in the past for tracking fast nanoscale dynamics,^10^ including SPT in living cells with up to 20 μs temporal resolution and about 15 nm spatial precision revealing signatures of hop diffusion.^11^

More recently, interferometric scattering microscopy (iSCAT)^12^ has demonstrated even higher spatiotemporal capabilities down to few nm precision^13^ and a few μs temporal resolution in vitro.^14^ Here, we use gold nanoparticles (AuNPs) coupled to GFP-tagged or YFP-tagged membrane proteins to demonstrate iSCAT-based SPT of membrane proteins with nanometer precision at kHz speeds in live epithelial cells and cultured hippocampal neurons. We use this approach to study if the increase in spatiotemporal precision allows for the detection of anomalous diffusion at the nanoscale. Such effects could arise from interactions between the plasma membrane and the cortical cytoskeleton, recently described in hippocampal neurons as a periodic actin-spectrin lattice.^15^

To achieve this, we functionalised 40 nm AuNPs with streptavidin and coupled them to biotinylated anti-GFP nanobodies. We then bound the functionalised nanoparticles to GFP- or YFP-tagged membrane proteins expressed in the U2OS human osteosarcoma cell line (Fig. 1a) or in cultured hippocampal rat neurons (Fig 1b). In both scenarios, the particles could be readily detected in unprocessed frames collected at 0.5 ms exposure times. Removing the background caused by diffracting material in cells (left and right panels) significantly improved the image quality, resulting in an average localisation error of σ = 3 nm, determined as the covariance-weighted mean square error of a 2D Gaussian fit, where σ_x_^2^ = C_x,x_ MSE, where C_x,x_ is the covariance of the variable x with itself. By constructing a trajectory from the localisations of a AuNP-labelled GPI-GFP protein on a U2SO cell at 2 kHz, we observe regions where the protein halts for up to 50 ms (100 frames) in regions a few tens of nm in diameter (Fig 1c, Supplementary movie 1). We observed some scattering fluctuations from cells, which varied in amplitude depending on the thickness and shape of the cell. On thin regions of U2OS cells and on neuron axons or dendrites we routinely observed background fluctuations of 1 – 2% root mean square (RMS). For comparison, the shot noise-induced background fluctuations under the current imaging conditions amount to approximately 0.3% RMS.^16^ The variations in background scattering caused by fluctuating cell material was thus the limiting factor preventing the use of smaller scattering labels if nanometer localisation precision was to be maintained.

**Fig 1:**
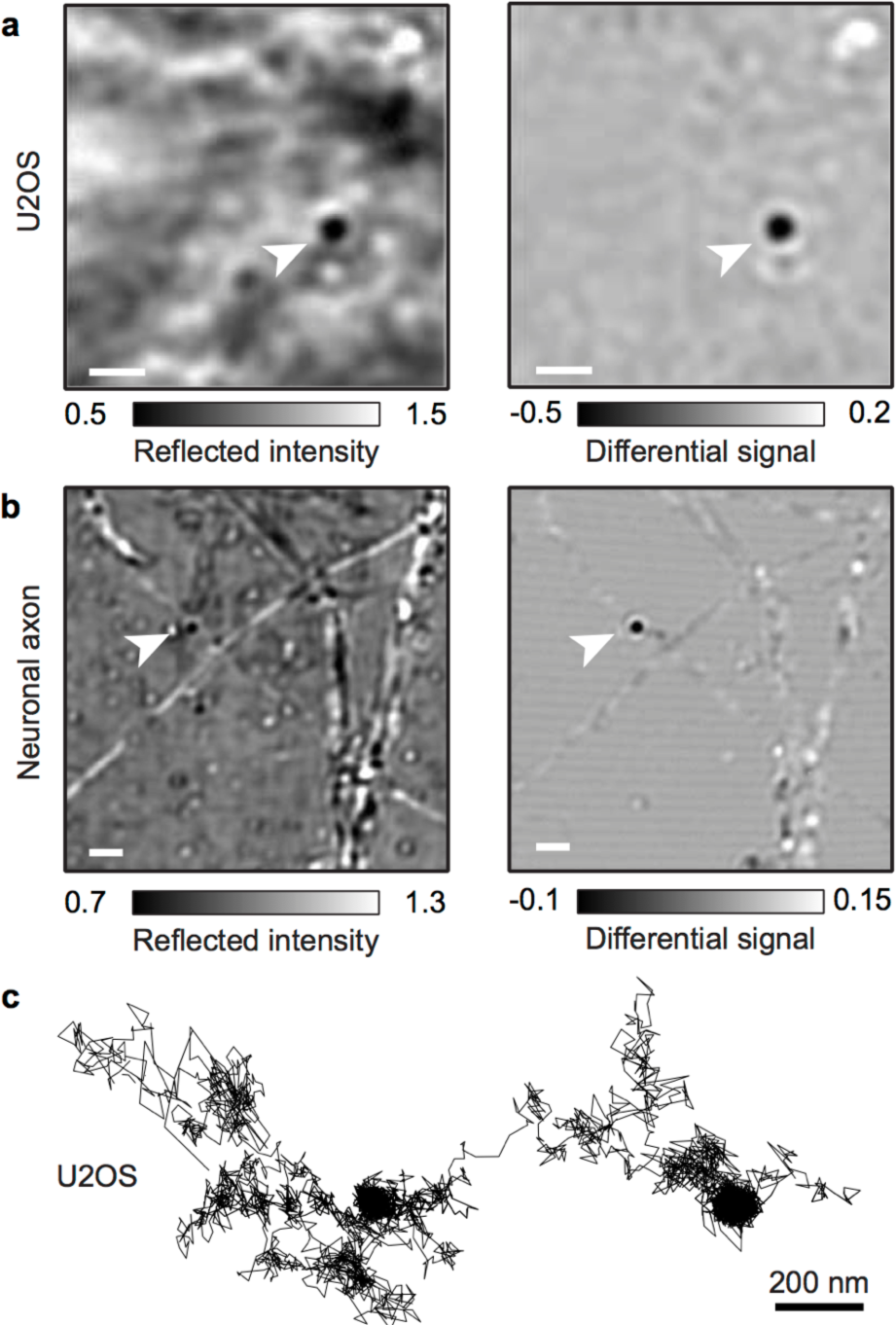
ISCAT images of a 40 nm AuNP-labelled membrane proteins on U2OS cells (a) and on neuronal cells (b). The gold particle is marked by an arrow in both the raw (left) and background subtracted (right) images. (c) Representative trajectory of a 40 nm AuNP-labelled protein in the U2OS cell membrane recorded at 2 kHz (10,000 frames total). Scale bars 2 μm.

We then performed SPT experiments at frame rates of 2 – 40 kHz in short bursts of 10’000 - 150’000 frames (3 – 20 s) on neurites of cultured hippocampal neurons (Fig 2). Neurons were imaged after day in vitro 6 (DIV 6) because by that point neuronal cells were well-developed and the axon extended far from the soma (Fig S2). This greatly increased the probability of finding a suitable region of the cell with a weakly fluctuating background. Trajectories for 40 nm AuNPs attached to membrane proteins were oriented along the long axis of the neurite (Fig 2a,b; Fig S3). When we determined the time-dependent one dimensional diffusion coefficients (D_1d_ - longitudinal motion) for our trajectories, we observed a significant reduction in D_1d_ for all membrane probes with increasing time lags (Fig 2c). This indicates that molecules exhibit subdiffusive motion on longer timescales, consistent with previous reports.^17^ We found subdiffusive behaviour for all probes tested: the transmembrane proteins L-YFP-GT46 (TM) and L-YFP-GTmHoneydew46 (TMX); and also for a GFP attached to the outer membrane leaflet via a glycosyl-phosphatidylinositol anchor (GPI-GFP). As shown before, the diffusion coefficient of GPI-GFP was about four times higher than for transmembrane proteins on timescales > 10 ms.^18^

**Fig 2:**
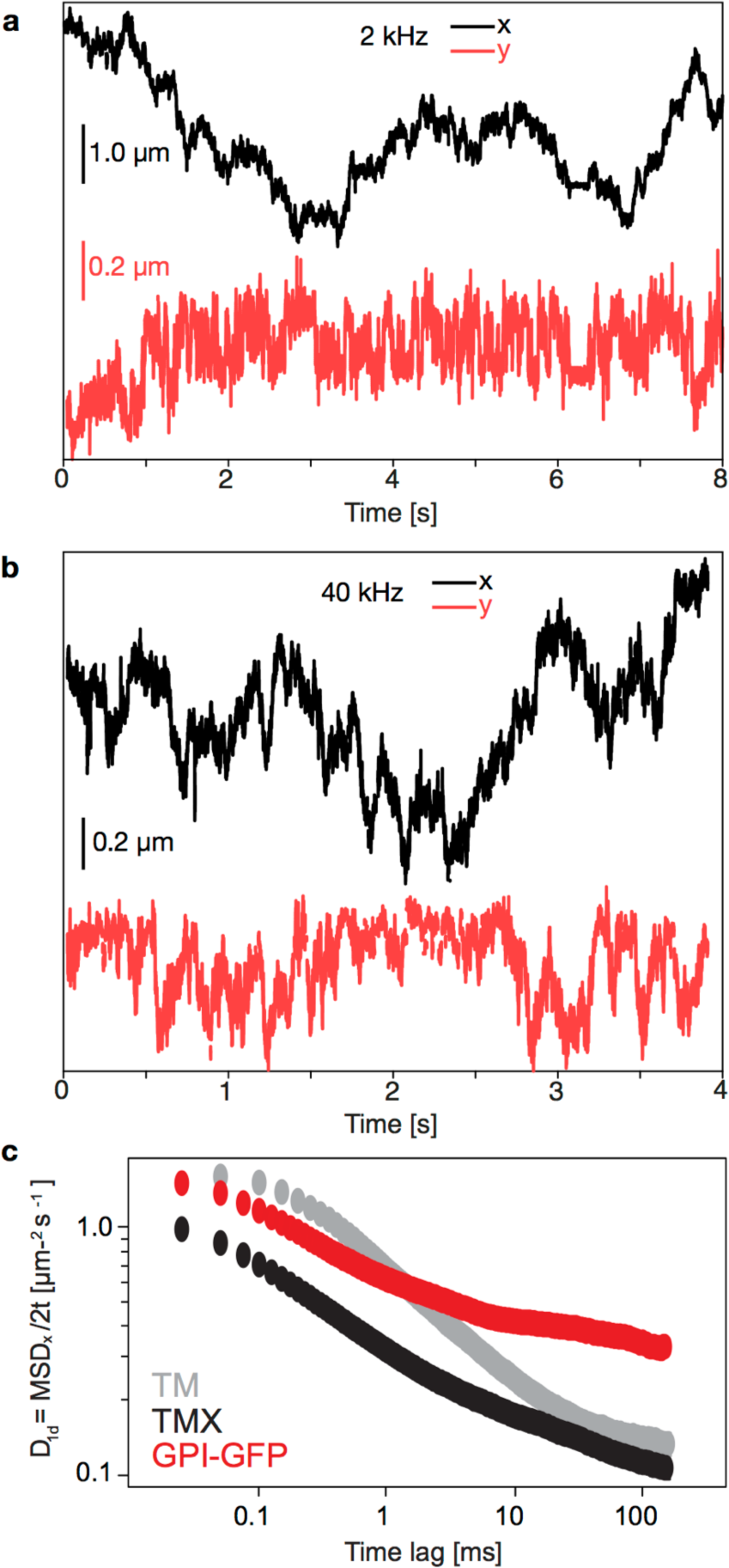
Analysis of 2D trajectories of 40 nm AuNP-labelled membrane protein probes. (a) Displacement in *x* (in direction of neurite propagation, black) and y (perpendicular to direction of neurite propagation, red) for a bead tracked at 2 kHz. (b) Same for bead tracked at 40 kHz. (c) Time-dependent 1D diffusion coefficient (x dimension) plotted against the time-lag used for analysis. Shown are plots for membrane proteins differing in their cytosolic domains and membrane anchors. TM = L-YFP-GT46, TMX = L-YFP-GTmHoneydew46, GPI-GFP = glycosylphosphatidylinositol anchored green fluorescent protein.

When we analyzed individual trajectories measured in neuronal cells, we did not observe transient periods of confinement in neurons as we did in U2OS cells (Fig 3, Fig S4). By visual examination, the iSCAT trajectories did not generally indicate the presence of compartments. On some, rare, occasions, however, a seemingly periodic array of areas of preferential residence of membrane molecules became apparent (Fig. 3a,b), similar to our previous observation of membrane compartments in the axon initial segment with QD-labelled membrane proteins.^19^ Numerous reports highlight the existence of a periodic cortical actin-spectrin network in axons^15,20-22^ and possibly dendrites ^23,24^, and we found the periodic arrangement of membrane compartments to be alternating with such actin rings in our fluorescence imaging studies.^19^ We thus asked whether we could find evidence for the same 180 – 200 nm periodicity in our iSCAT trajectory data. We applied a moving window of 5 ms over the frame-to-frame displacement of the paricles and plotted the 20% of positions with the lowest standard deviation over this window in blue (Fig 3a). Such statistical analyses preferentially detect weak confinement events, which are overshadowed by random motion (see also Fig S5). For these datapoints, a repetitive pattern was clearly visible from the plotted positions in several trajectories. On the other hand, when we plotted the 20% of positions that showed the lowest confinement in this analysis (yellow), we found that no such pattern was detectable. The repetitive pattern seen in the autocorrelation analysis of confined localisations followed a 180 – 200 nm periodicity (Fig 3b), comparable to that of the of the actin-spectrin cytoskeleton in neurites. This observation agrees with our previous description of a periodic pattern in localisations from SPT with quantum dots (QDs) on live neurons.^19^ We remark that we here only tracked single AuNPs over few seconds, compared to multiple QDs being tracked over several min in our previous work, demonstrating the high sensitivity of the iSCAT approach, enabled by the comparatively high amount of datapoints retrievable from single trajectories. Compared to U2OS cells, neurons presented additional challenges as the diameter of a neurite is typically 200 – 500 nm with an approximately circular shape and therefore high curvature.

**Fig 3:**
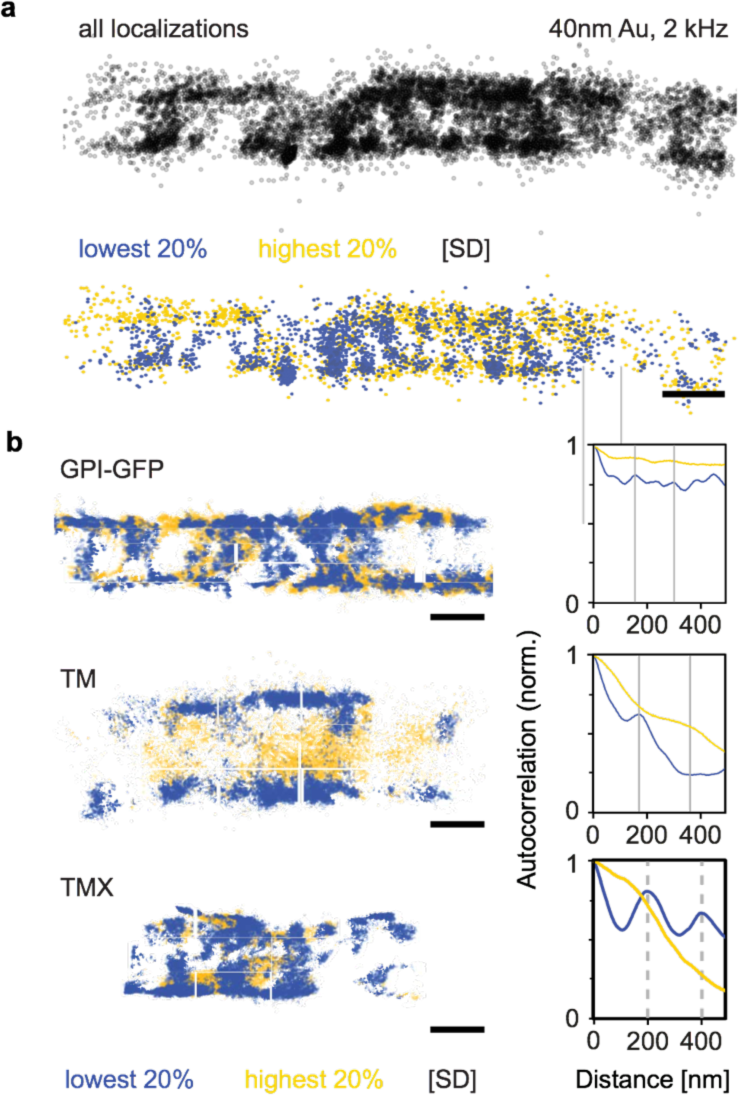
Analysis of membrane protein mobility on neurites. (a) Analysis of 20’000 datapoints from a 2 kHz tracking experiment of a 40 nm AuNP-labelled GPI-GFP protein (black, top). The 20% of data points exhibiting the lowest confinement are shown in yellow, the 20% with highest confinement are shown in blue (below). Low/high confinement are classified as high/low standard deviation of displacement over a 5 ms moving window. (b, left) Confinement analysis of different membrane proteins in neurites. TM = L-YFP-GT46, TMX = L-YFP-GTmHoneydew46, GPI-GFP = glycosylphosphatidylinositol anchored green fluorescent protein. (b, right) Autocorrelations along the propagation direction of the neurite of the most confined (blue) and least confined (yellow) 20% of datapoints from the dataset. Scale bars 200 nm.

Owing to the interferometric nature of iSCAT, the contrast of AuNPs varied with its height above the coverglass (Fig 4a). For 40 nm AuNPs, the contrast ranged from -50% to +50%, yielding a maximum SNR of 50%/2% = 25 at 635 nm. We took advantage of this observation and assigned a z-value to our 2D SPT data based on changes in contrast (Fig S7) to render our localisations in 3D (Fig 4b). We note that the density in the *y* coordinate is less amenable to interpretation for several reasons: the projection of the cross section of the neurite onto the *y* axis produces a higher density of localisations at the circumference of the neurite. Furthermore, the localisation density in *y* depends on the proximity to the zero-contrast plane, the gold particle being undetectable in this plane. We remark that the three-dimensional information is qualitative, since a precise calibration of iSCAT signal vs particle height above the coverglass is challenging in this experimental arrangement.^25^

**Fig 4:**
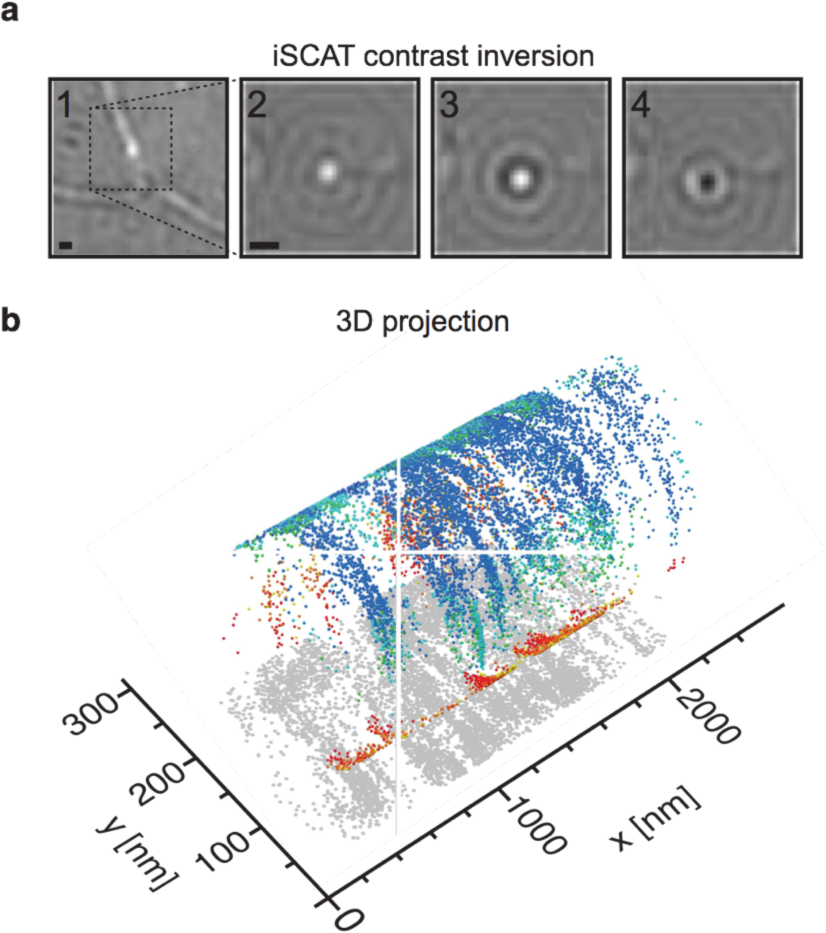
3D iSCAT tracking on live neurons. (a) Individual frames from a movie of a 40 nm AuNP diffusing on a neurite, exhiting variations in contrast as a function of time. (b) A 3D trajectory on a neurite reconstructed from the x,y-coordinates, using the circular geometry *y*^2^ + *z*^2^ = *r*^2^ (Fig S7). The sign of the *z*-coordinate was determined by applying the condition that contrast varies continuously with *z*. Contrast values are color-coded from red to blue. The recorded 2D data is displayed in grey, projected onto the *x/y* plane. Scale bars 500 nm.

We have shown here that iSCAT SPT with a spatial resolution of few nm and a temporal resolution in the microsecond range is possible in live cells. Using this approach, we found that the apparent diffusion coefficient of membrane molecules indeed depends on the timelag with which the mean-square displacement is measured and calculated as proposed previously.^11^ This has been interpreted as indicative of membrane compartmentalisation,^17^ since according to the Singer-Nicolson model, diffusion should be free in the plasma membrane.^26^ Kusumi et *al.* suggested a model in which membrane protein diffusion is free on the microsecond and nm scale, but confined in domains of ~10s to 100s of nms in diameter in the plasma membrane in dependence on the submembrane actin cytoskeleton.^27^ This notion is supported by recent observations in our laboratory revealing confinement of GPI-GFP motion between 200 nm spaced actin rings in the axon initial segment.^19^ Using iSCAT, we did not observe clearly defined compartments, yet still find signatures of subdiffusive motion on the tens of ms time scale. Spatially-resolved analysis of protein motion suggests a periodicity consistent with actin ring spacings, although we cannot confirm axon initial segment localisation. Measurements close to the soma where the axon initial segment is located were impeded by a highly fluctuating background and a low probability of encountering AuNPs.

It will be exciting to investigate in future work if membrane compartmentalisation can be correlated to specific locations in neurons. Furthermore, the basis for subdiffusive motion on longer timescales could be transmembrane interaction with other submembrane agents. In previous experiments in supported lipid bilayers, we showed that the motion of crosslinked lipids in the top bilayer leaflet can be influenced by the immobilisation of lipids in the lower bilayer leaflet.^13^ Future work aimed at identifying the specific factors that control transmembrane confinement of membrane molecules in cells will be an important contribution to the understanding of the dynamics and organisation of the plasma membrane. The high spatiotemporal precision of iSCAT together with the potential for three-dimensional tracking provides a potentially powerful, complementary approach to fluorescence-based methodologies in live cells.

